# H3K4 methylation dependent and independent chromatin regulation by *JHD2* and *SET1* in budding yeast

**DOI:** 10.1101/265926

**Authors:** Kwan Yin Lee, Ziyan Chen, River Jiang, Marc D. Meneghini

## Abstract

Set1 and Jhd2 regulate the methylation state of histone H3 lysine-4 (H3K4me) through their opposing methyltransferase and demethylase activities in the budding yeast *Saccharomyces cerevisiae*. H3K4me associates with actively transcribed genes and, like both *SET1* and *JHD2* themselves, is known to regulate gene expression diversely. It remains unclear, however, if Set1 and Jhd2 act solely through H3K4me. Relevantly, Set1 methylates lysine residues in the kinetochore protein Dam1 while genetic studies of the *S. pombe SET1* ortholog suggest the existence of non-H3K4 Set1 targets relevant to gene regulation. We interrogated genetic interactions of *JHD2* and *SET1* with essential genes involved in varied aspects of the transcription cycle. Our findings implicate *JHD2* in genetic inhibition of the histone chaperone complexes Spt16-Pob3 (FACT) and Spt6-Spn1. This targeted screen also revealed that *JHD2* inhibits the Nrd1-Nab3-Sen1 (NNS) transcription termination complex. We find that while Jhd2’s impact on these transcription regulatory complexes likely acts via H3K4me, Set1 governs the roles of FACT and NNS through opposing H3K4-dependent and -independent functions. We also identify diametrically opposing consequences for mutation of H3K4 to alanine or arginine, illuminating that caution must be taken in interpreting histone mutation studies. Unlike FACT and NNS, detailed genetic studies suggest an H3K4me-centric mode of Spt6-Spn1 regulation by *JHD2* and *SET1*. Chromatin immunoprecipitation and transcript quantification experiments show that Jhd2 opposes the positioning of a Spt6-deposited nucleosome near the transcription start site of *SER3*, a Spt6-Spn1 regulated gene, leading to hyper-induction of *SER3*. In addition to confirming and extending an emerging role for Jhd2 in the control of nucleosome occupancy near transcription start sites, our findings suggest some of the chromatin regulatory functions of Set1 are independent of H3K4 methylation.

## Introduction

Methylation of histone H3 lysine-4 (H3K4me) represents one of the most comprehensively studied chromatin modifications. Like most histone lysine methylations, H3K4 exhibits mono, di, and tri-methylated states (H3K4me1, H3K4me2, and H3K4me3) that have distinctive regulatory outputs reflecting the recruitment of effector proteins (SANTOS-ROSA *et al.* 2003; TAVERNA *et al.* 2006; KIM AND BURATOWSKI 2009). In the budding yeast *Saccharomyces cerevisiae,* H3K4me levels are controlled by the opposing functions of two highly conserved enzymes belonging to the Trithorax/MLL and JARID1 families: the Set1 methyltransferase and the Jhd2 demethylase (KROGAN *et al.* 2002; INGVARSDOTTIR *et al.* 2007; LIANG *et al.* 2007).

Unless otherwise stated, whenever we refer to *SET1* and *JHD2*, we are signifying the budding yeast genes specifically. While Jhd2 appears to act alone, Set1, like Trithorax/MLL, functions within COMPASS, a conserved complex of proteins that function in Set1 targeting and regulation (KROGAN *et al.* 2002). Surprisingly, despite the crucial role of Trithorax/MLL and JARID during development and disease in animals, neither *SET1* nor *JHD2* are essential for cell viability in budding yeast (BENEVOLENSKAYA *et al.* 2005; DEY *et al.* 2008; LOPEZ-BIGAS *et al.* 2008). Moreover, *JHD2* null mutants (*jhd2*Δ) have surprisingly limited phenotypic impact in otherwise wild-type (WT) mitotic cells grown in the presence of glucose, confounding the study of this important chromatin regulatory protein using yeast (LENSTRA *et al.* 2011; XU *et al.* 2012).

Previously, we found that *JHD2* exhibits specific functions during the gametogenesis phase of the budding yeast life cycle, also known as sporulation (XU *et al.* 2012). The formation of robust spores requires that *JHD2* maintain a period of productive transcription in the face of encroaching and developmentally programmed transcriptional quiescence. During this period, *JHD2* globally demethylates H3K4 at intergenic regions upstream of transcription start sites (TSSs) and overlapping with transcription termination sites (TTSs). Associated with these H3K4me defects, *JHD2* also represses the accumulation of noncoding intergenic transcripts genome-wide (XU *et al.* 2012). These findings suggest roles for *JHD2* in diverse aspects of transcription, including those related to termination and its associated mRNA processing pathways. Indeed, a recent study found that Jhd2 physically interacts with CPF (cleavage and polyadenylation factor) and causes defects in mRNA 3’ untranslated region length for some genes (BLAIR *et al.* 2016). More recently, we determined that H3K4 demethylation by Jhd2 occurs not only during sporulation, but also in mitotic cells in response to non-fermentable carbon sources or to nitrogen manipulations that lead to increased levels of alpha-ketoglutarate, an intermediary metabolite essential for demethylase activity of Jhd2 and the entire Jumonji family of demethylases to which JARIDs belong (LIANG *et al.* 2007; SOLOVEYCHIK *et al.* 2016).

Our previous studies identified developmental/nutritional contexts during which *JHD2* impacts gene expression, but the mechanisms remain opaque. To gain further insight into genetic pathways by which *JHD2* and *SET1* impact gene expression, we used a targeted screening approach. Because genome-wide screens have been unsuccessful in the identification of gene deletions that exhibit interactions with *jhd2Δ* ((COSTANZO *et al.* 2016), our unpublished findings), we selected essential components of the transcriptional machinery and tested whether temperature sensitive (TS) mutants of these components genetically interact with *jhd2*Δ, and in most cases, *set1*Δ as well. Using this approach, we find that *JHD2* genetically inhibits the essential RNA PolII transcription regulatory complexes Spt6-Spn1, FACT (facilitates chromatin transcription), and NNS (Nab3-Nrd1-Sen1) (FORMOSA *et al.* 2001; VASILJEVA AND BURATOWSKI 2006; MCDONALD *et al.* 2010). More detailed genetic experiments reveal that *JHD2*’s inhibitory role can be attributed solely to reversal of H3K4me. Accordingly, genetic interrogation of *SET1* confirms a positive regulatory impact of H3K4me, but also illuminates a counterbalancing and inhibitory role of *SET1* that acts independent of H3K4me. Using molecular biological experiments, we determine that Jhd2 opposes the positioning of a Spt6-Spn1-deposited nucleosome near the 3’ of the *SRG1* non-coding transcript, and accordingly impacts the induction of the downstream *SER3* gene known to be regulated by *SRG1*, perhaps explaining at least some of the basis for the genetic interactions. In summary, our findings identify three transcriptional regulatory complexes subject to regulation by Jhd2 and Set1, and illuminate that *SET1* impinges upon them through H3K4me dependent and independent means.

## Results

### Genetic interactions of *JHD2* with essential transcription regulatory factors

We used a targeted screening approach to identify mutants that exhibit genetic interactions with *jhd2*Δ. Because Jhd2 reverses a histone modification associated with active transcription, we evaluated temperature sensitive (TS) alleles of essential genes related to transcriptional control as interacting candidates of *jhd2*Δ. We used strains sourced from a library created and curated previously (LI *et al.* 2011), with the only exceptions being *nab3-42* and *spt16-319*, which we obtained separately (O’DONNELL *et al.* 2004; DARBY *et al.* 2012). All strains described here were engineered using genetic crosses and tetrad dissection. Strain fitness was assessed using spot assays to compare growth rates at varied temperatures. Deletion of *JHD2* or *SET1* did not cause growth defects under these conditions (Fig. 1A, B, and C). For all results shown, we isolated at least 2 independently constructed strain replicates through tetrad dissection. Though replicates and WT control strains are not always shown in the interest of space, all results we report here are upheld in these replicates.

**Figure 1.**
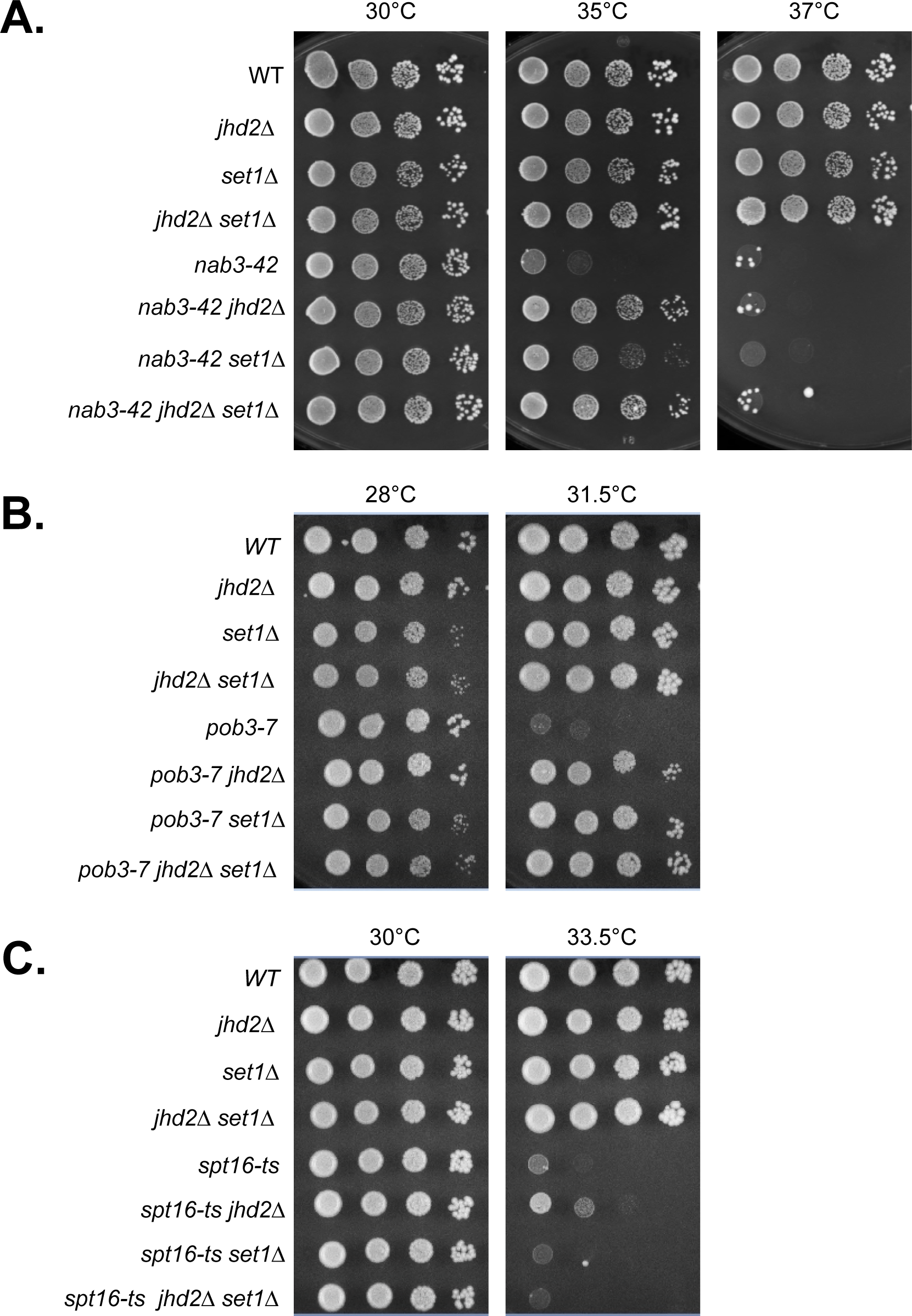
TS alleles of NNS and FACT subunits exhibit genetic interactions with H3K4 modifying enzymes. Yeast strains with the indicated genotypes were serially diluted ten-fold, spotted onto agar plates containing synthetic complete media, and grown at indicated temperatures. Genetic interactions of *jhd2*Δ and *set1*Δ with different temperature sensitive alleles of NNS and FACT subunits are shown: (A) *nab3-42* (B) *pob3-7* (C) *spt16-ts*.

Given the physical association of Jhd2 with CPF/CF, we first evaluated *jhd2*Δ genetic interactions with TS mutations in 10 different subunits of this large complex (BLAIR *et al.* 2016). Somewhat surprisingly, we failed to identify any reproducible interactions with CPF/CF (Table 1). NNS functions alongside CPF as one of two distinct mRNA processing pathways in budding yeast. Nab3 and Nrd1 encode RRM domain containing proteins that recognize specific sequences in nascent transcripts (STEINMETZ AND BROW 1996; CARROLL *et al.* 2004; CREAMER *et al.* 2011). The Sen1 subunit is homologous to the human helicase protein Senataxin and provides the activity that physically dislodges PolII from the DNA (PORRUA AND LIBRI 2013; CHEN *et al.* 2014). While the majority of mRNA 3’ end processing of protein coding genes occurs through CPF, NNS controls the transcriptional termination and processing of noncoding RNAs. The most widely studied targets of NNS are snRNAs, snoRNAs, and cryptic untranslated transcripts (CUTS) (URSIC *et al.* 1997; STEINMETZ *et al.* 2001; ARIGO *et al.* 2006; THIEBAUT *et al.* 2006).

**Table 1.** Genetic interactions of *JHD2* with essential transcription regulatory factors. Specific TS allele is listed in the left column the associated protein complex of each factor shown in the corresponding middle column. In the right column, any observed genetic interactions with *jhd2*Δ are documented. N.I. no interaction, + positive interaction (+ weakest, ++ medium, +++ strong suppression), - weak negative interaction.

We found that *jhd2*Δ robustly suppressed 2 different alleles of *NAB3*, *nab3-42* and *nab3-11* (Table 1, Fig. 1A and S1A). Deletion of *JHD2* also modestly suppressed *sen1-1* (Table 1, Fig. S1B). As shown, incubation at high temperature was sufficient to eliminate growth of *nab3-42*, confirming that *jhd2*Δ did not provide bypass suppression and that Jhd2 therefore acted to inhibit *nab3-42* function at semi-permissive temperatures (Fig. 1A). Though we only show the lack of bypass suppression for *nab3-42* in the interest of space, this was the case with all interactions we report here. NNS termination is coupled with polyadenylation by the TRAMP complex and RNA processing by Exosome (VASILJEVA AND BURATOWSKI 2006; VANACOVA AND STEFL 2007). We observed a similar suppressive interaction of *jhd2*Δ with the sole essential TRAMP component, *MTR4* (Table 1 and data not shown). Interestingly, the only exosome subunit encoding gene we investigated, *MTR3*, provided the solitary example of a negative interaction with *JHD2*; *jhd2*Δ caused a modest but reproducible enhancement of the *mtr3-ts* growth defect (Table 1 and data not shown).

Among the numerous other essential genes we interrogated, including those contributing to the functions of mediator, RNAPII and its associated C-terminal domain modifications, and TATA binding protein (TBP), only temperature sensitive alleles of genes encoding Spt6-Spn1 and FACT subunits exhibited genetic interactions with *jhd2*Δ (Table 1, Fig. 1B and 1C). Spt6-Spn1 and FACT are conserved heterodimeric histone chaperone complexes that enable RNA PolII transcription through chromatin, deposit nucleosomes in the wake of elongating polymerase, and suppress spurious intragenic transcription (KAPLAN *et al.* 2003; DUINA 2011; HAINER *et al.* 2011a). While Spt6-Spn1 and FACT have many common and overlapping functions, they exhibit differences in histone binding specificity and in their abilities to disrupt nucleosomes during transcription (BORTVIN AND WINSTON 1996; ORPHANIDES *et al.* 1999; BELOTSERKOVSKAYA *et al.* 2003; MCCULLOUGH *et al.* 2015). FACT can disrupt and reassemble nucleosomes, while Spt6-Spn1 only has the ability to reassemble them (ORPHANIDES *et al.* 1998; ORPHANIDES *et al.* 1999; BELOTSERKOVSKAYA *et al.* 2003). We also confirmed a previous finding that the temperature sensitive growth defect of *ess1-H146R*, which encodes a CTD-associated prolyl-isomerase, was suppressed by *jhd2*Δ (MA *et al*. 2012).

### H3K4me dependent and independent modulation of NNS and FACT by *JHD2* and *SET1*

As *JHD2* reverses H3K4me, we hypothesized that increased H3K4me caused by *jhd2*Δ accounted for the genetic interactions described above. Indeed, H3K4me3 has been suggested to promote NNS function, with *set1*Δ enhancing the growth and termination phenotypes caused by alleles of *NRD1* that delete portions of its N-terminus (TERZI *et al.* 2011). Terzi et al. also showed that *set1*Δ enhanced termination defects caused by *nab3-11*, though they did not evaluate *SET1*’s impact on *nab3-11* growth properties (TERZI *et al.* 2011). We therefore tested if *jhd2*Δ suppression of NNS mutations could occur in the absence of *SET1.* Unexpectedly, *set1*Δ suppressed the growth defect of *nab3-42* to a degree that exceeded *jhd2*Δ suppression, with the triple mutants resembling *nab3-42 set1*Δ (Fig. 1A). TERZI *et al.* 2011 found that H3K4me3 promotes NNS function using truncation alleles of *NRD1* (TERZI *et al*. 2011), but the suppression of *nab3-42* by *set1*Δ presented here suggested that H3K4me antagonizes NNS function. To rule out some unusual allele specific interaction of *nab3-42* with *set1*Δ, we replicated the *set1*Δ and *jhd2*Δ interactions with *NAB3* using the *nab3-11* allele (Fig. S1A). Furthermore, and consistent with the *NAB3* results, *sen1-1* was also suppressed by both *jhd2*Δ and *set1*Δ, again with the triple mutants resembling *sen1-1 set1*Δ (Fig. S1B). While none of our experiments overlap with those of Terzi et al, their findings would not have predicted ours nor vice versa. In particular, our finding that *set1*Δ suppressed the *nab3-11* growth defect is somewhat discordant with Terzi et al’s finding of an aggravating consequence of *set1*Δ on *nab3-11* for transcriptional termination. Among the many possible explanations for this discrepancy, a simple view is that the essential functions of *NAB3* may not be directly reflected by termination defects described by Terzi et al.

Our targeted screen also identified the FACT subunit encoding alleles *pob3-7* and *spt16-ts* as exhibiting alleviating genetic interactions with *jhd2*Δ (Table 1, Fig. 1B and 1C). Using the same experimental approach as with NNS subunits, we tested if *jhd2*Δ suppression of *pob3-7* and *spt16-ts* required *SET1*. Curiously, and unlike the case with NNS subunits, *pob3-7* and *spt16-ts* exhibited distinct genetic interactions in combination with *set1*Δ, even though both factors function in the FACT complex. We found that *set1*Δ suppressed the growth defect of *pob3-7* mutants to a level comparable to the suppression conferred by *jhd2*Δ, with the triple mutants showing no evidence of synergy (Fig. 1B). In contrast, the *spt16-ts* temperature sensitive growth defect was suppressed by *jhd2*Δ in a *SET1* dependent manner, suggesting that H3K4me promotes Spt16 function (Fig. 1C). We confirmed these genetic interactions with additional alleles of *POB3* and *SPT16*, *pob3-L78R* and *spt16-319* (Fig. S1C and S1D). The differences in *SET1* genetic interaction phenotypes observed for *SPT16* and *POB3* mutants suggest complexities in nucleosome interactions, or that their encoded proteins may exhibit separable functions apart from the FACT complex.

To investigate how both *set1*Δ and *jhd2*Δ can rescue certain temperature sensitive growth defects, we employed strains that express H3K4 amino acid substitution mutants from a chromosomal locus in place of wild type H3 (DAI *et al.* 2008). The most widely utilized amino acid substitutions for this approach are alanine (H3K4A) and arginine (H3K4R). We first confirmed the histone mutations by sequencing and then determined that H3K4A and H3K4R alone or in combination with *set1*Δ did not cause any growth phenotypes under the conditions of our spotting assays (Fig. S2A). Unexpectedly, we found that H3K4A and H3K4R modified *pob3-7* and *nab3-11* mutants in opposing fashion. H3K4A suppressed *pob3-7* and *nab3-11* temperature sensitivity (Fig. 2A and S2B). Suppression by H3K4A and *set1*Δ did not display any synergy. H3K4R caused the opposite effect of H3K4A, exacerbating the temperature sensitive growth defect of *nab3-11* and *pob3-7* (Fig. 2A, 2B, 2C, S2B, and S3A). We note that this latter result is consistent with previous findings suggesting a positive impact for H3K4me3 on NNS function (TERZI *et al.* 2011).

**Figure 2.**
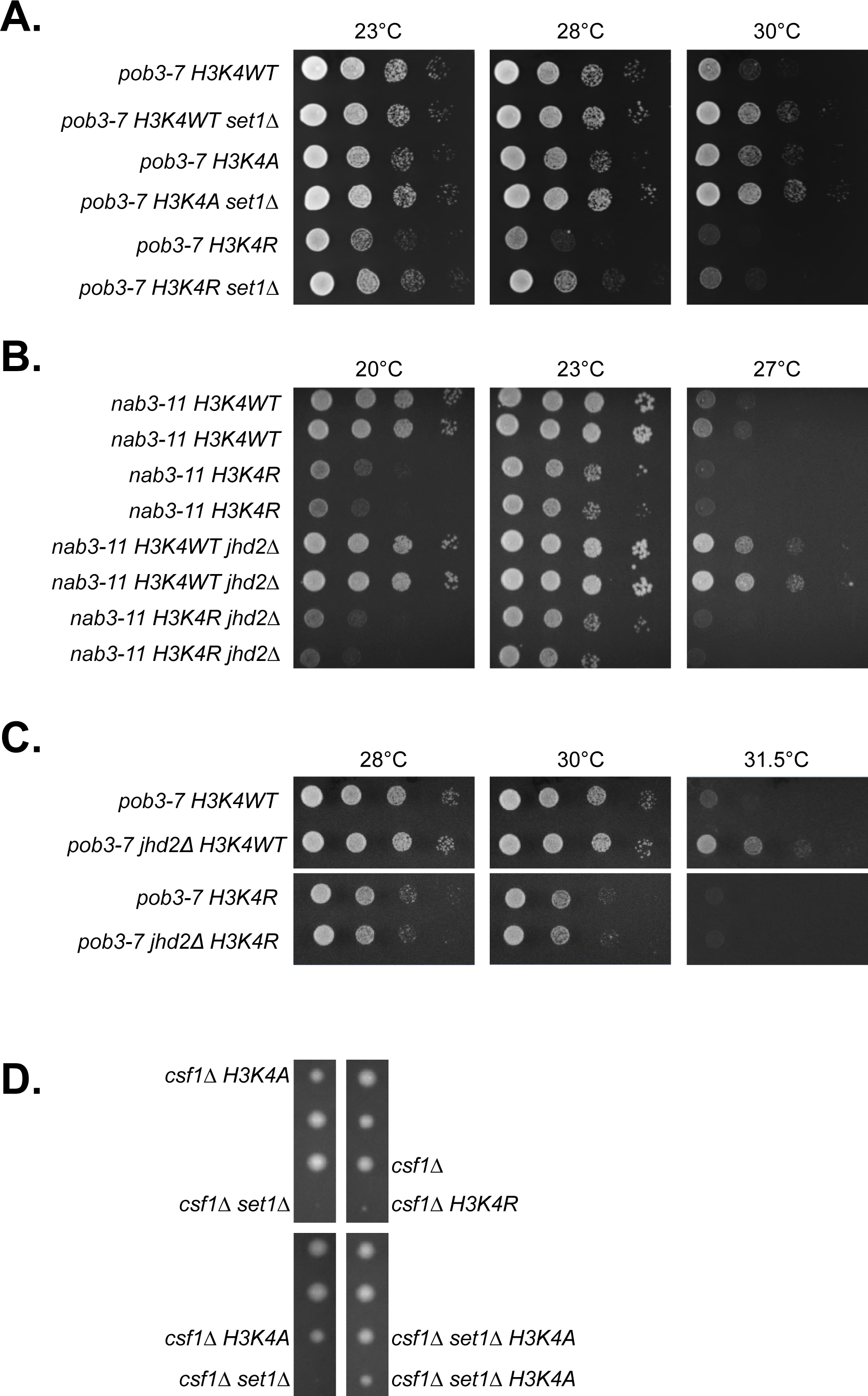
Genetic suppression of NNS and FACT by *set1*Δ and *jhd2*Δ dependent and independent of H3K4me. Plate spot assays (as described in Figure 1) were used to compare the growth of the indicated strains. Isogenic strains were constructed through genetic crosses with H3K4A and H3K4R substitution mutants from the Dharmacon Non Essential Histone H3 & H4 Mutant Collection. (A) Genetic interactions of *set1*Δ, H3K4A, and H3K4R with *pob3-7*. (B) Genetic interactions of *jhd2*Δ and H3K4R with *nab3-11*. Growth of two independent isolates of each genotype is shown. (C) Genetic interactions of *jhd2*Δ and H3K4R with *pob3-7*. (D) Diploid strains containing *csf1*Δ, *set1*Δ, and either H3K4A or H3K4R were sporulated and dissected onto YPD medium. A representative field of the haploid ascospores recovered and their respective genotypes are shown.

To shed light on the confounding differences in H3K4A and H3K4R phenotypes, we performed additional studies exploiting an annotated synthetic lethal interaction between *set1*Δ and *csf1*Δ (COSTANZO *et al.* 2016). *CSF1* (cold sensitive for fermentation) encodes a conserved yet poorly characterized protein required for growth on glucose at low temperatures and is involved in secretory protein maturation and amino acid metabolism. (TOKAI *et al.* 2000; COPIC *et al.* 2009; COSTANZO *et al.* 2016). In support of the interpretation that *csf1Δ set1*Δ lethality was caused by loss of H3K4me, we found that *jhd2*Δ suppressed *csf1*Δ and that *JHD2* overexpression using an integrated galactose-inducible allele enhanced *csf1*Δ (Fig. S2C and S2D). The *csf1*Δ enhancement by *GAL-JHD2* was abrogated by mutation of histidine-427 to alanine (H427A), which renders Jhd2 catalytically inactive (INGVARSDOTTIR *et al.* 2007; LIANG *et al.* 2007) (Fig. S2C and S2D).

We constructed diploid strains heterozygous for *csf1*Δ, *set1*Δ, and H3K4A or H3K4R mutants and interrogated them using tetrad dissections. In addition to confirming *csf1Δ set1*Δ synthetic lethality, we found that H3K4R exhibited synthetic lethality with *csf1*Δ (Fig. 2D). In stark contrast, H3K4A had no consequence for *csf1*Δ viability, and remarkably, suppressed the synthetic lethality of *csf1Δ set1*Δ (Fig. 2D). We dissected dozens of tetrads and confirmed that this pattern of inheritance was highly stereotypical and not due to random spore inviability (Table 2). It is noteworthy that while both H3K4A and H3K4R abolish H3K4 methylation, arginine retains the positively charged characteristic of lysine in contrast to the aliphatic alanine residue. Accordingly, arginine much more closely resembles the structure of a lysine residue compared with alanine. These results suggest that H3K4R more reliably recapitulates the consequences of *set1*Δ for chromatin structure and point toward neomorphic characteristics of H3K4A, which seem likely to manifest differently depending on genetic context. We therefore proceeded with our studies using the H3K4R mutant.

**Table 2.** Observed and expected genetic frequencies of *csf1*Δ with histone H3K4A and H3K4R amino acid substitution mutants. Diploid strains mmy7658 (H3K4R), mmy7659 (H3K4WT), and mmy7660 (H3K4A) were sporulated and dissected onto YPD medium. These strains are heterozygous for *set1*Δ, *csf1*Δ, and the corresponding histone amino acid substitution. The observed frequency of each genotype was calculated by dividing to the total number of spores counted and is shown in the parentheses.

To reconcile the H3K4R mutant phenotype with suppression of NNS and FACT subunits by *set1*Δ, we constructed strains combining H3K4WT or H3K4R with *set1*Δ and NNS/FACT mutants. We found that *set1*Δ conferred suppression of the growth phenotypes of *pob3-* 7, *nab3-11,* and *sen1-1* even in the context of H3K4R albeit not to the extent seen in H3K4WT (Fig. 2A, S2B, S3A, and S3B). Thus, the suppression of NNS and FACT subunits by *set1*Δ in H3K4R mutants suggests that some activity of Set1 acted in opposition to NNS and FACT independent of H3K4me (Fig. 3).

**Figure 3.**
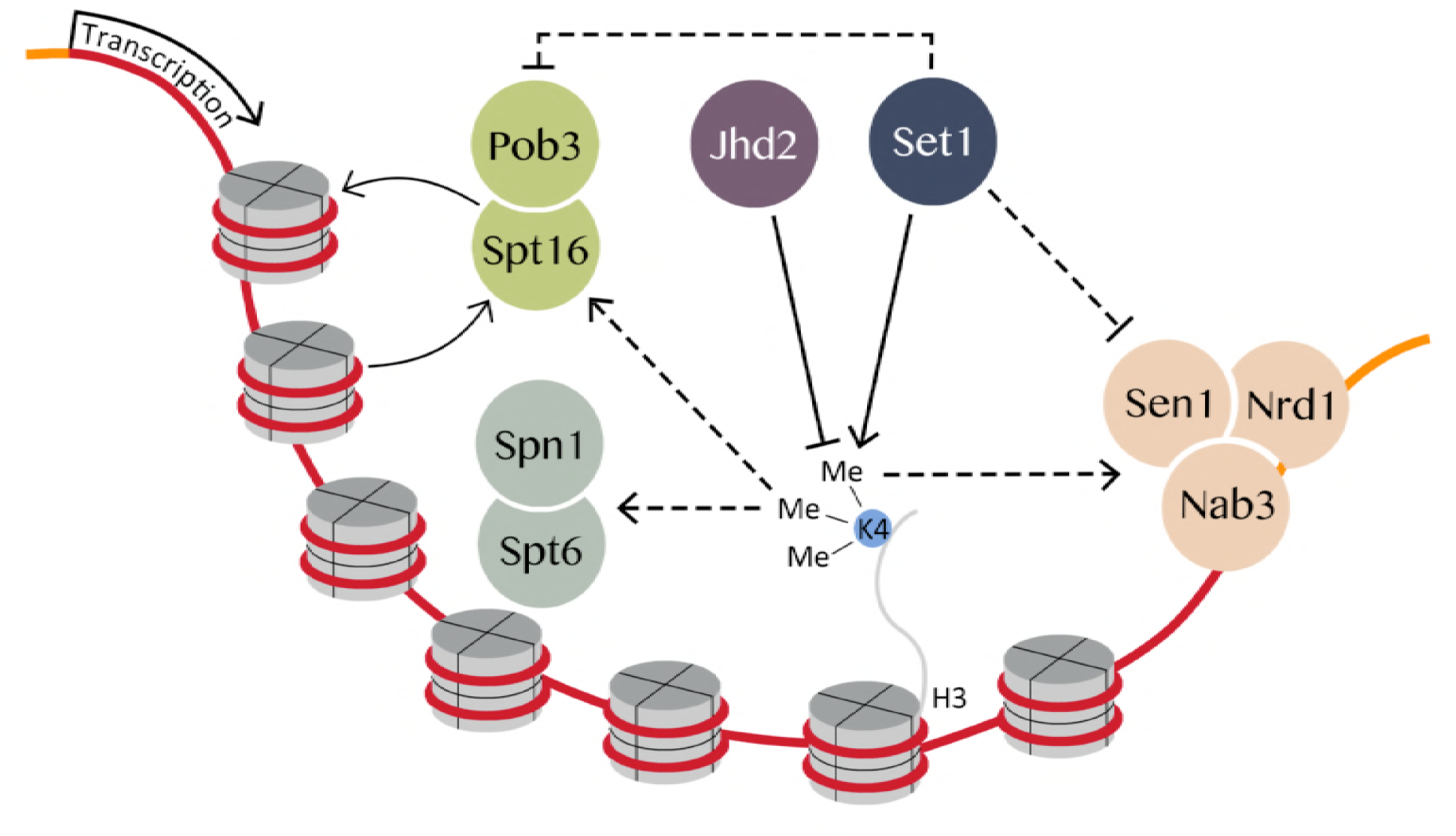
Genetic model of FACT and NNS regulation by Set1 and Jhd2. Methylation of H3K4 promoted FACT and NNS function while Set1 exhibited an inhibitory function independent of H3K4. Spt6-Spn1 is positively regulated by H3K4me3.

To further evaluate the positive impact of H3K4me on NNS and FACT and the role of *JHD2* in this, we performed analogous experiments with *jhd2*Δ. Suppression of *nab3-11*, *pob3-7*, and *sen1-1* by *jhd2*Δ could no longer occur in the context of H3K4R, arguing that Jhd2 opposed NNS and FACT function solely through reversal of H3K4me (Fig. 2B, 2C, and S3C). Based on these results as well as previous findings from Terzi et. al, we propose counterbalancing regulation of NNS and FACT by Set1 and Jhd2, with H3K4me promoting their functions and an unknown inhibitory impact conferred by Set1 in a H3K4 independent manner (Fig. 3).

To gain insight into how Set1 opposed FACT independent of H3K4, we constructed strains that had the endogenous *SET1* locus replaced with *SET1-G951S*, an allele that abolishes Set1 enzymatic activity, but does not disrupt the integrity of COMPASS (NAGY *et al.* 2002; SCHIBLER *et al.* 2016). We found that *SET1-G951S* suppressed *pob3-7* to an extent that was equivalent to *set1*Δ in strains expressing a WT allele of H3K4 (Fig. S4). As enzymatically dead alleles of *SET1* lead to a depletion of Set1 protein levels, this was to be expected (SOARES *et al.* 2014). Curiously, in strains expressing H3K4R, which do not deplete Set1 levels to nearly the extent seen with enzymatically dead alleles of Set1 (SOARES *et al.* 2014), *SET1-G951S* failed to suppress *pob3-7* at 30° C, but did suppress equivalently to *set1*Δ at the more stringent temperature of 31.5 °C (Fig. S4). Similar results were obtained with *nab3-11* (data not shown). These results, while complex, suggest that the H3K4-independent inhibition of *pob3-7* by Set1 may involve both enzymatic and non-enzymatic roles of this protein (Fig. 3).

### Spt6-Spn1 was opposingly governed by *JHD2* and *SET1* through H3K4me3

The complexity of interactions exhibited by NNS and FACT with *SET1* rendered them difficult to further study without a more detailed understanding of what regulatory functions *SET1* has independent of H3K4. We therefore extended our genetic analysis to Spt6-Spn1. As mentioned above, we determined that TS alleles of both Spt6-Spn1 subunits, *spt6-14* and *spn1-K192N*, were suppressed by *jhd2*Δ (Table 1, Fig. 4A and 4B). In contrast to NNS and FACT, the temperature sensitive growth defects of both Spt6-Spn1 subunits were suppressed by *jhd2*Δ in a *SET1* and H3K4 methylation dependent manner (Fig. 4A, B, C, and data not shown). Moreover, and consistent with the interpretation that *SET1*-mediated H3K4me promoted Spt6-Spn1 function, both *set1*Δ and H3K4R enhanced the growth defects of *spt6-14* and *spn1-K192N* (Fig. 4A, B, C, and data not shown).

**Figure 4.**
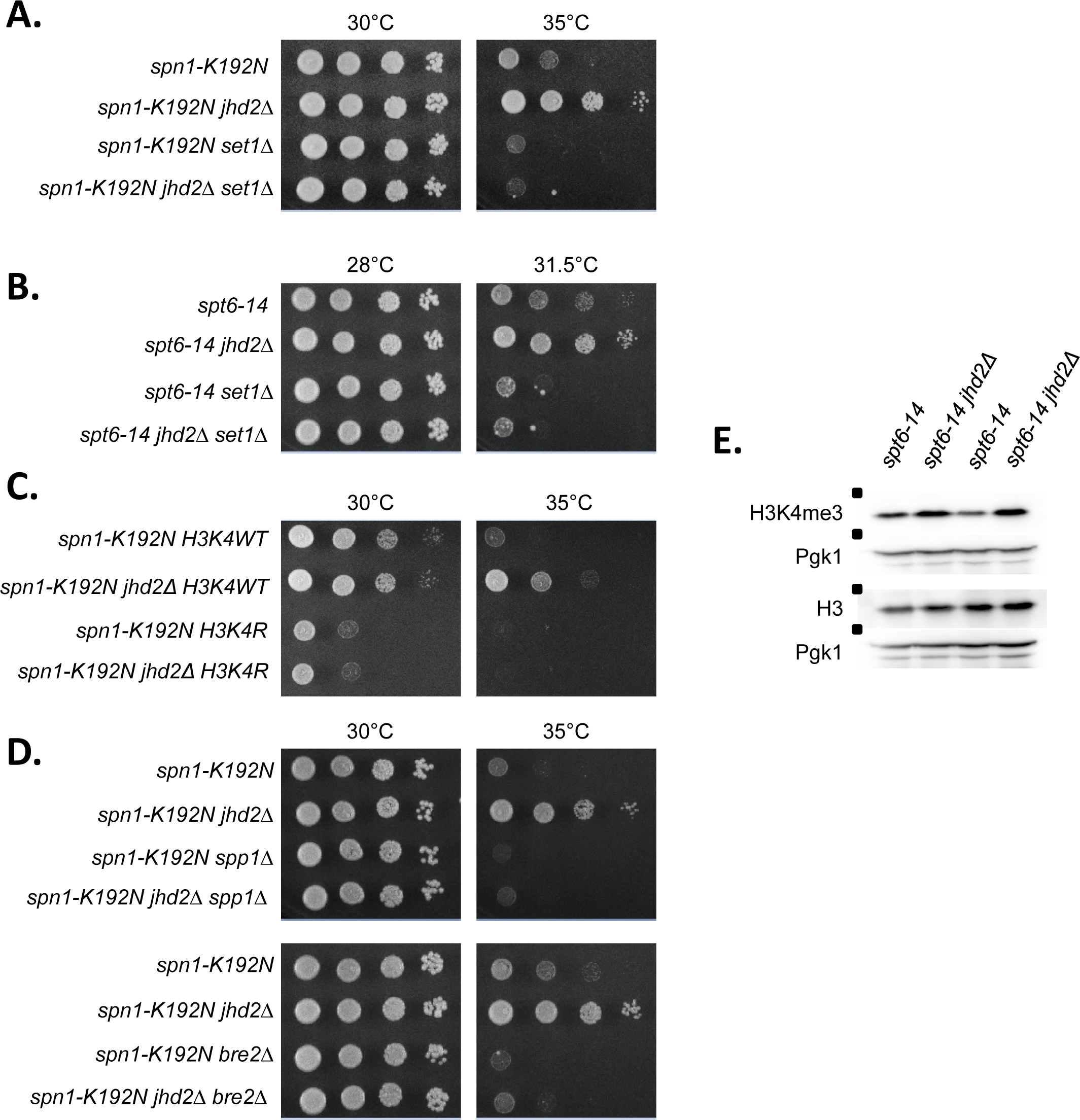
Spt6-Spn1 complex temperature sensitive mutants are rescued by an increase in H3K4me3 through *jhd2*Δ. Plate spot assays (as described in Figure 1) were used to compare the growth of the indicated strains. Genetic interactions of *jhd2*Δ and *set1*Δ with Spt6-Spn1 are shown: (A) *spt6-14* (B) *spn1-K192N*. (C) Isogenic Strains were constructed through genetic crosses with the H3K4R substitution mutant from the Dharmacon Non Essential Histone H3 & H4 Mutant Collection. Genetic interactions of *jhd2*Δ and H3K4R with *spn1-K192N*. (D) Genetic interactions of *jhd2*Δ, nonessential COMPASS subunits *spp1*Δ and *bre2*Δ, and *spn1-K192N*. (E) Western blot analysis of H3K4me3, Histone H3, and Pgk1 were performed on biological duplicates of *spt6-14* and s *pt6-14 jhd2*Δ. Pgk1 serves as a loading control.

To validate that Jhd2 demethylation of H3K4 opposed Spt6-Spn1, we complemented the *jhd2*Δ-suppressed strains with previously described *JHD2* expressing plasmids (MERSMAN *et al.* 2009). Cells harboring these plasmids express *JHD2* under the control of a constitutive *PYK1* promoter, which leads to modestly increased levels of Jhd2 and associated decreased H3K4me3 levels. These effects on H3K4me could be abrogated by the H427A mutation or by deletion of its conserved PHD domain, which may enable Jhd2 recruitment to chromatin (MERSMAN *et al.* 2009). As expected, we found that *jhd2*Δ suppression of *spt6-14* and *spn1-K192N* could be reverted by *JHD2* complementation and that the growth defects of *spt6-14* and *spn1-K192N* were in fact enhanced by the *JHD2* overexpressing plasmids (Fig. S5A and S5B). Importantly, this suppression was lost in the H427A and PHD domain mutants (which both encode stable proteins (MERSMAN *et al.* 2009)) (Fig. S5A and S5B). Our findings support the conclusion that suppression of Spt6-Spn1 alleles by *jhd2*Δ was due to the loss of H3K4 demethylation.

To gain insights into which H3K4 methylation species accounted for suppression of Spt6-Spn1, we genetically interrogated COMPASS complex subunits known to specifically perturb H3K4me3 or H3K4me3 and me2. The effects of *bre2*Δ and *spp1*Δ on H3K4me have been extensively described. Deletion of *BRE2* results in a complete loss of H3K4me3 and a significant decrease of H3K4me2 but no change in H3K4me1, while *spp1*Δ results in a substantial decrease of H3K4me3 with no detectable consequence on bulk H3K4me2 or me1 (SCHNEIDER *et al.* 2005; DEHE *et al.* 2006; TAKAHASHI *et al.* 2009; MARGARITIS *et al.* 2012). As with *set1*Δ, we found that both *bre2*Δ and *spp1*Δ enhanced the growth defect of *spn1-K192N* (Fig. 4D). Moreover, *jhd2*Δ suppression of *spn1-K192N* was completely dependent on *BRE2* and *SPP1* (Fig. 4D). We compared bulk H3K4me3 levels by western blot in *spt6-14* and *spt6-14 jhd2*Δ at a semi-permissive temperature. Consistent with the genetic data, we observed a modest increase in bulk H3K4me3 levels in the *spt6-14 jhd2*Δ double mutant (Figure 4E). We confirmed the reproducibility of this by quantifying the relative abundance of H3K4me3/H3 as well as of H3/Pgk1 from three biological replicates of *spt6-14* and *spt6-14 jhd2*Δ (Fig. S6). Collectively, our data support the conclusion that *JHD2* and *SET1* opposingly governed the function of Spt6-Spn1 through H3K4me3 (Fig. 3).

### *JHD2* repressed TSS-associated nucleosome occupancy and transcriptional induction of a Spt6-Spn1 regulated gene

Reflecting its role in the recycling of nucleosomes in the wake of elongating PolII, Spt6-Spn1 mutation causes reduced histone occupancy genome-wide (IVANOVSKA *et al.* 2011). We asked if *jhd2*Δ suppression of Spt6-Spn1 involved a reversal of these histone occupancy defects. First, we used western blots to measure bulk histone H3 levels and found that *jhd2Δ, set1Δ,* and *spt6-14* mutants showed no apparent defects in histone H3 levels at the permissive temperature of 23°C (Figure 5A). At the semi-permissive temperature of 30°C *spt6-14* exhibited a dramatic reduction in H3 levels, consistent with its role as a histone chaperone (Figure 5A). This reduction was not reversed by *jhd2*Δ suggesting that suppression did not occur by generically improving Spt6-Spn1 histone recycling or chaperone function (Figure 4E, 5A, and S6).

**Figure 5.**
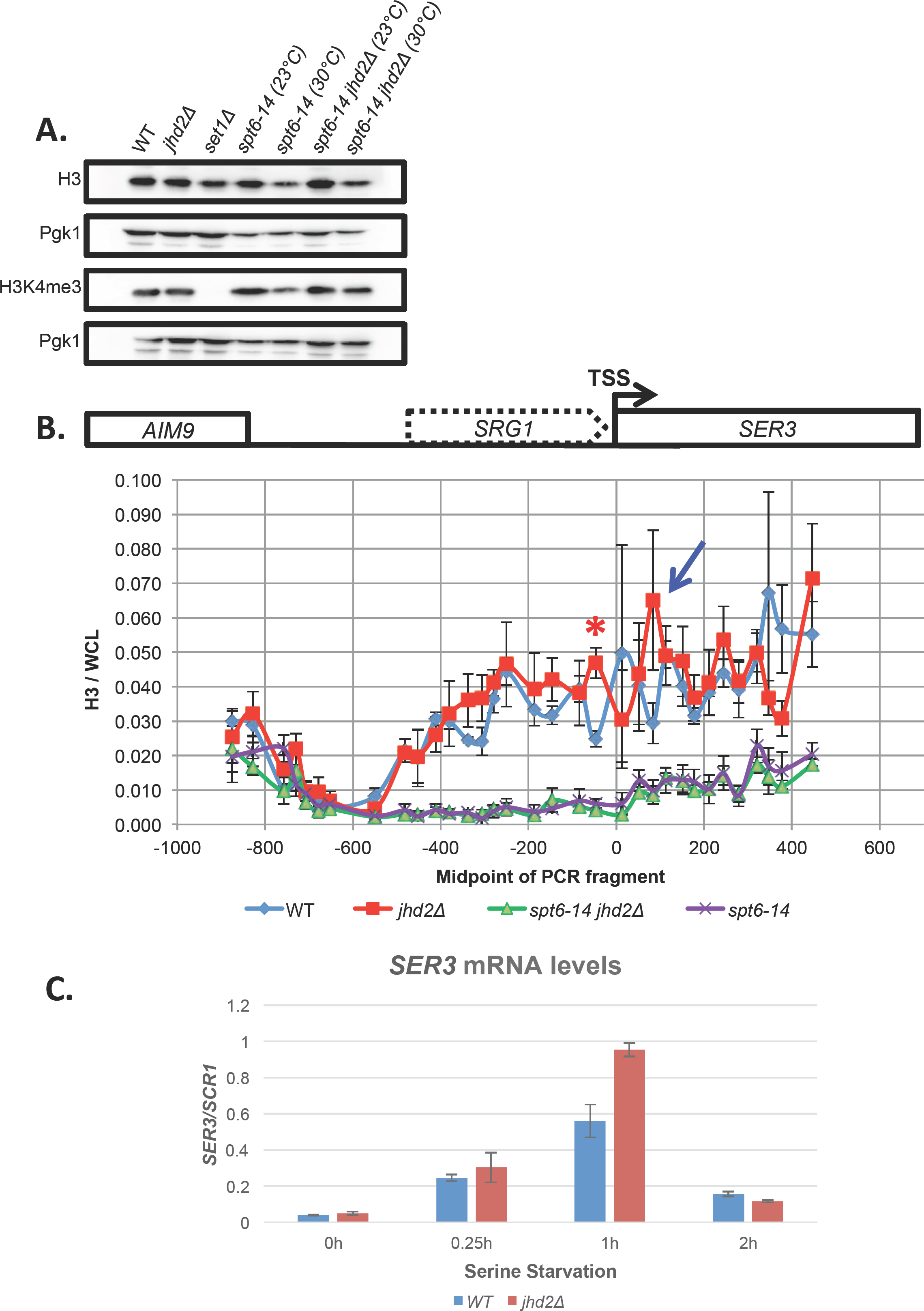
*JHD2* deletion does not rescue Spt6-Spn1 mutants by compensating for defects in chromatin structure. (A) Western blot detection of H3K4me3, pan-H3, and Pgk1 are shown from extracts of wildtype and mutant cells grown at 23°C (permissive) and 30°C (semi-permissive temperature). Pgk1 serves as a loading control. (B) Histone H3 ChIP was performed on strains grown at 30°C (semi-permissive temperature) across the *SRG1-SER3* locus (schematic of locus shown above graph). H3 ChIP signal was normalized to signal from whole cell lysate (WCL) DNA. The transcription start site of *SER3* is positioned at 0 and for *SRG1* at -480. Error bars represent standard error of the mean. Significance as calculated by a two-tailed student’s t-test is shown where \**p* < 0.05. A blue arrow denotes an additional H3 ChIP peak in *jhd2*Δ that was not statistically significant. (C) *SER3* transcript level was measured by RT-qPCR at specific intervals after serine withdrawal in wildtype and *jhd2*Δ mutants. *SER3* signal was normalized to *SCR1.*

To gain more detailed insight into the relationship between Jhd2 and Spt6-Spn1, we measured chromatin-associated histone H3 in WT, *jhd2Δ, spt6-14,* and *spt6-14 jhd2*Δ cells at the *SRG1-SER3* locus using chromatin immunoprecipitation with an anti-H3 antibody followed by real time PCR quantification of the immunoprecipitated DNA (ChIP-qPCR). In order to evaluate nucleosome positions at high resolution across *SRG1-SER3*, we used 33 closely spaced PCR amplicons to measure immunoprecipitated DNA (HAINER *et al.* 2011b). The averaged results of 3 independent biological experiments are shown (Fig. 5B). As previously described, an abundance of H3 was detected throughout the *SRG1* transcript unit extending into the *SER3* gene in WT cells. These H3 levels were dramatically depleted in *spt6-14* mutants grown at the semi-permissive temperature, confirming that their deposition was Spt6 dependent (HAINER *et al.* 2011b) (Fig. 5B). Consistent with our western blotting results, *spt6-14 jhd2*Δ mutants did not exhibit improved H3 deposition at *SRG1-SER3* at the semi-permissive temperature we used (Fig. 5B). In WT and *jhd2*Δ cells with normal *SPT6* function, the H3 ChIP profiles were nearly superimposable throughout the entire *SRG1-SER3* region with one conspicuous exception: *jhd2*Δ caused a highly significant increase in H3 abundance at a single distinctive peak near the 3’ end of the *SRG1* transcript unit, presumably reflecting a well-positioned nucleosome (Fig. 5B, peak denoted with a red asterisk, *p* < 0.05). This *JHD2*-repressed nucleosome was positioned ∼75 base pairs upstream of the *SER3* TSS, and positioned ∼75 base pairs downstream of this TSS we observed an additional H3 ChIP peak in *jhd2*Δ, though this second peak was not statistically significant (Fig. 5B, denoted with a blue arrow). Spanning the *SER3* TSS, H3 ChIP signal was noisy in both WT and *jhd2Δ,* suggesting a dynamic state of nucleosomal occupancy at this region (Fig. 5B).

We performed our ChIP experiments from cells grown in rich media, a condition in which *SER3* transcription is repressed in a manner dependent on Spt6-Spn1-deposited nucleosomes across the *SER3* TSS in response to *SRG1* transcription (PRUNESKI AND MARTENS 2011) (Fig. 5B). A simple encapsulation of our findings posits that *JHD2* prevented Spt6-Spn1-deposited nucleosomes from accumulating at positions on the flanks of the *SER3* TSS, resulting in dynamic and noisy nucleosome occupancy across the *SER3* TSS (Fig. 5B). We detected no differences in *SER3* abundance from WT and *jhd2*Δ cells grown in rich media, suggesting that this altered chromatin architecture did not impact *SER3* repression (data not shown). To determine if it instead impacted *SER3* induction, we subjected WT and *jhd2*Δ cells to serine starvation and compared their relative *SER3* mRNA abundance using quantitative PCR amplification of reverse transcribed RNA (RT-qPCR). We found that *SER3* was maximally expressed 1 hour after serine withdrawal, and that *SER3* mRNA accumulated to ∼1.7-fold increased levels in *jhd2*Δ at this timepoint (Fig 5C). Following 2 hours of serine starvation, yeast cells adapt and repress *SER3* induction, and *jhd2*Δ had no consequence for this repression (MARTENS *et al.* 2005) (Fig. 5C). Thus, the phased nucleosomes flanking the *SER3* TSS we observed in *jhd2*Δ were associated with a hyper-induction of *SER3*.

## Discussion

In this work, we identify novel genetic interactions between the H3K4 demethylase *JHD2* and genes encoding subunits of the essential transcription regulatory complexes NNS, FACT, and Spt6-Spn1. Our genetic findings support the conclusion that reversal of H3K4me by Jhd2 opposes the functions of these complexes (Fig. 3). In the case of NNS, our findings are in good agreement with a previous study suggesting that H3K4me3 promotes NNS function (TERZI *et al.* 2011). A simple prediction from our findings is that *set1*Δ, which abolishes all H3K4 methylation, should cause the opposite phenotype as, and exhibit epistasis to, *jhd2*Δ. While this was indeed the case for Spt6-Spn1 and the *SPT16* subunit of FACT, TS alleles of *SEN1*, *NAB3*, and *POB3* were each suppressed by *set1*Δ to an extent equivalent to, or exceeding that by *jhd2*Δ. More detailed genetic studies revealed that *SET1* opposed the functions of *SEN1*, *NAB3*, and *POB3* in a manner that was independent of H3K4me. We point out that the Swd2 subunit of COMPASS also functions within the NNS-related termination complex APT, which could potentially explain the different NNS interactions with *SET1* deletion versus K4 mutation shown in this study (Nedea et al., 2003; Soares and Buratowski, 2012). Our genetic model posits that H3K4me positively influences NNS and FACT functions, and that Set1 exhibits an additional counterbalancing H3K4me-independent activity that negatively impacts these complexes (Fig. 3). For Spt6-Spn1, our genetic data suggested a positive impact by H3K4me3 with no repressive H3K4-indpendent role for Set1 (Fig. 3). Our molecular studies showed that Jhd2, and by inference H3K4me reversal, negatively regulated the accumulation of Spt6-Spn1 deposited nucleosomes flanking the *SER3* TSS, resulting in hyper-induction of *SER3* upon serine starvation.

In contrast to our findings and what has been reported previously (HAINER *et al.* 2011b), Ramakrishnan *et al* found that both *jhd2*Δ and *set1*Δ caused dramatic up-regulation of *SER3* in rich serine-replete conditions (RAMAKRISHNAN *et al.* 2016). Under such conditions *SER3* is normally strongly repressed, and we found that levels were so low as to be nearly undetectable with no difference between WT and *jhd2*Δ (Fig. 5C) (MARTENS *et al.* 2004; MARTENS *et al.* 2005; HAINER *et al.* 2011b). To reconcile these discrepancies, we point out that all of the experiments in the Ramakrishnan *et al* study were performed in a strain background that utilized plasmid expressed histone H2A-H2B, with the endogenous H2A-H2B genes deleted, a condition known to reduce histone dosage and alter transcription (CLARK-ADAMS *et al.* 1988). This is not to say we reject the findings of Ramakrishnan *et al;* rather, the findings of Ramakrishnan *et al* seem to illuminate a facile way to investigate chromatin biology in yeast, by ‘sensitizing’ the chromatin through reduced histone dosage. Whatever the case may be, our findings uphold an emerging role for Jhd2 in the control of nucleosome occupancy at TSS’s and TTS’s. *JHD2*’s known developmental role in opposing nucleosome accumulation at these regions may be attributable to it antagonistic impact on Spt6-Spn1 (XU *et al.* 2012). As Spt6-Spn1, Set1, Jhd2, and H3K4me are conserved throughout eukaryotes, these mechanisms may have broad significance.

An important feature of our findings relates to functions of Set1 that are independent H3K4 methylation. The only confirmed non-H3K4 enzymatic target of Set1, or indeed of any member of this highly conserved family of enzymes, is the essential kinetochore protein Dam1. Dam1 lysine residue methylation by Set1 regulates the phosphorylation status of neighboring serine residues by mediating the opposing activities of Ipl1 and Glc7 kinase and phosphatase (ZHANG *et al.* 2005). *Schizosaccharomyces pombe* Set1 (_sp_Set1) controls transcription at silent mating loci, telomeric regions, and retrotransposon (Tf) elements, as well as the clustering of diverse genomic regions encoding Tf elements. Interestingly, _sp_Set1 accomplishes these sundry functions in a manner dependent on differing domains of Set1 that do, or do not, have consequences for H3K4me. The two key regulatory roles of _sp_Set1 appear dependent on _sp_Set1’s enzymatic activity and on its RNA binding RRM2 domain, a domain found in both fission and budding yeast Set1 (TRESAUGUES *et al.* 2006; LORENZ *et al.* 2014; MIKHEYEVA *et al.* 2014). Consistent with the RRM2 having important regulatory significance independent of H3K4me, a recent study identified many mRNA targets bound by the budding yeast Set1 RRM2 domain (SAYOU *et al.* 2017). Thus, the RRM2 domain may underlie at least some of the H3K4me-independent regulatory functions of Set1 we identify here. Our genetic results, however, also suggest that enzymatic activity of Set1 directed towards a non-H3K4 substrate has regulatory significance, as may be the case for *S. pombe* Set1 (LORENZ *et al.* 2014; MIKHEYEVA *et al.* 2014). It seems highly plausible that chromatin-associated factors may be targets of regulatory lysine methylation by Set1 and other histone methyltransferases.

## Materials and Methods

### Yeast Strains and Plasmids

Standard yeast genetic methods were used for construction of all strains. Yeast strains and plasmids used in this study are listed in Supplementary Data Table 1. All strains were constructed through genetic crosses followed by dissections the BY4742 background. Yeast strains were inoculated into several mL of YPD (1% yeast extract, 2% peptone, and 2% glucose) and grown overnight at room temperature (23°C). Each strain was diluted to an OD_600_ = 0.4, serially diluted five times and spotted on synthetic complete media (YNB media (Multicell Wisent) containing 5 g/L of ammonium sulfate and either 2% glucose or 2% galactose.

### Western blots

Exponential cultures with OD_600_ between 1-2 were lysed by vortexing with acid washed glass beads in SUMEB buffer as described previously (Fredrickson et al. 2011). Total protein concentration was quantified using an RC/DC assay (BioRad). Equal amounts of protein were electrophoresed on 12% SDS-PAGE gels, and transferred onto Amersham Hybond-P membranes (GE). Immunoblot analysis was performed using standard procedures. All blots were scanned with a ChemiDoc XRS+ Imaging System. Band intensities were quantified using ImageJ 1.51 v software.

### Chromatin immunoprecipitation

Strains were grown at 30°C in YPD to OD_600_ 0.8. Crosslinking was performed in 1% formaldehyde for 15 minutes at room temperature (23°C) and then immediately quenched with glycine at a final concentration of 125 mM. Cells were pelleted and washed 2 times with ice cold PBS. Cell pellets were snap frozen in 2 mL screwcap tubes using liquid nitrogen and stored at -80°C. Frozen pellets were resuspended in lysis buffer (50 mM HEPES pH 7.5, 140 mM NaCl, 1 mM EDTA pH 8.3, 1% Triton X-100, 0.1% NaDOC) without thawing and topped off with glass beads. Cells were lysed through multiple rounds of beadbeating and then sonicated to shear the chromatin to fragment sizes of approximately 200 to 500 bp. Cross-linked chromatin fragments were immunoprecipitated with antibody overnight. Protein A-Sepharose beads were then added to the samples, and samples were incubated for 90 minutes. The immunoprecipitated complexes were washed with lysis buffer, lysis buffer containing 500 mM NaCl, wash buffer (10 mM Tris-HCl pH 8.0, 0.25M LiCl, 0.5% NP40, 0.5% NaDOC, 1mM EDTA pH 8.3), and TE (10 mM Tris-HCl pH 8.0, 1 mM EDTA). Next, the immunoprecipitated chromatin was eluted from beads with elution buffer (50 mM Tris-HCl pH 8.0, 10 mM EDTA, 1% SDS) and then eluted with TE/0.67% SDS (10 mM Tris-HCl pH 8.0, 1 mM EDTA pH 8.3, 0.67% SDS, 0.334 mg/mL proteinase K). The two elutions were combined. Whole cell extract samples were topped off with TE/1% SDS (10 mM Tris-HCl pH 8.0, 1mM EDTA, pH 8.3, 1% SDS, 0.25 mg/mL proteinase K). Formaldehyde cross-linking was reversed by incubating the eluates at 65°C overnight. DNA from the eluates was treated with RNAse A and purified with a QIAquick PCR purification kit (Qiagen). Immunoprecipitated fractions (IP) and whole-cell extracts (Input) containing DNA were analyzed by real-time PCR as described previously (XU *et al.* 2012) using the primers listed in Supplementary Data Table 2. SYBR green signal was measured using the BioRad iQ5 Multicolor Real Time PCR Detection System.

### *SER3* RT-qPCR analysis

5-10 OD equivalents were harvested for RNA extraction. For RT-qPCR, RNA was extracted with acidic phenol at 65°C for 30 min. RNA was then purified, precipitated and resuspended in RNAse free water. cDNA was prepared as described previously using random priming (XU *et al.* 2012). Quantitative PCR quantification of cDNA using SYBR green was measured using the BioRad iQ5 Multicolor Real Time PCR Detection System as described previously (XU *et al.* 2012). Primer sequences for *SER3* and *SCR1* are shown in Supplementary Data Table 2. Transcript levels for each primer pair tested were normalized to the reference transcript *SCR1*.

### *SER3* induction assays

Yeast strains were inoculated into several mL of YPD and grown overnight at room temperature (23°C). Each strain was diluted to an OD_600_ = 0.4 in synthetic complete media and grown to OD_600_ = 2 at 30°C. At this time, cells were harvested by centrifugation, washed with water, and resuspended in synthetic media lacking serine. Growth of cells was resumed with constant shaking at 30°C. Cell samples were collected at specific intervals and frozen using liquid nitrogen for analysis by RT-qPCR as described previously (XU *et al.* 2012). Primer sequences for *SER3* and *SCR1* are shown in Supplementary Data Table 2. RNA isolation, cDNA synthesis, and qPCR is described above.

## Acknowledgements

This work was supported by CIHR grant MOP-89996 to M.D.M. We are particularly grateful to Dr. Charlie Boone and Dr. Brenda Andrews for access to their temperature-sensitive mutant library and other reagents. We would like to thank Dr. Jeffry L. Corden for the *nab3-42* mutant. We thank Dr. Scott D. Briggs for *PYK1-JHD2*, *PYK1-JHD2(H427A),* and *PYK1-JHD2(PHD*Δ) plasmids. The *SET1-G951S* and *spt16-319* mutant strains were kindly provided by Dr. Sharon Dent and Dr. Richard A. Singer.

---

**Supp. Fig. 1.** Alternative NNS and FACT subunit TS alleles exhibit the same genetic interaction profile as *NAB3*.

Yeast strains with the indicated genotypes were serially diluted ten-fold, spotted onto agar plates containing synthetic complete media, and grown at indicated temperatures. Genetic interactions of *jhd2*Δ and *set1*Δ with different temperature sensitive alleles of NNS and FACT subunits are shown: (A) *nab3-11* (B) *sen1-1* (C) *pob3-L78R* (D) *spt16-319*.

**Supp. Fig. 2.**Arginine substitution of H3K4 most reliably mimics *set1Δ.*

Plate spot assays (as described in previous figures) were used to compare the growth of the indicated strains. (A) Isogenic Strains were constructed through genetic crosses with H3K4A and H3K4R substitution mutants from the Dharmacon Non Essential Histone H3 & H4 Mutant Collection. Genetic interactions of H3K4A and H3K4R with *set1*Δ. (B) Genetic interactions of *set1*Δ and H3K4A with *nab3-11*. (C) Genetic interaction of *jhd2*Δ and *csf1*Δ is shown. (D) Yeast strains with the indicated genotypes were serially diluted ten-fold, spotted onto agar plates containing synthetic complete media containing glucose or galactose to compare the growth of the indicated strains. *JHD2* is expressed under the control of a galactose inducible promoter. The H427A mutation disrupts the histone demethylase activity of Jhd2.

**Supp Fig. 3.** Genetic suppression of NNS by *set1*Δ and *jhd2*Δ dependent and independent of H3K4me.

Plate spot assays (as described in previous figures) were used to compare the growth of the indicated strains. Isogenic Strains were constructed through genetic crosses with H3K4A and H3K4R substitution mutants from the Dharmacon Non Essential Histone H3 & H4 Mutant Collection. (A) Genetic interactions of *set1*Δ and H3K4R with *nab3-11*. Growth of two independent isolates of each genotype is shown. (B) Genetic interactions of *set1*Δ and H3K4R with *sen1-1.* (C) Genetic interactions of *jhd2*Δ and H3K4R with *sen1-1.*

**Supp Fig. 4.**H3K4-independent suppression of *pob3-7* by Set1 may involve both enzymatic and non-enzymatic activities.

Plate spot assays (as described in previous figures) were used to compare the growth of the indicated strains. Analyzed is a G951S mutation that disrupts histone methyltransferase activity of Set1.

**Supp Fig. 5.**The enzymatic activity and the PHD domain of Jhd2 opposes Spt6-Spn1.

*JHD2, JHD2(H427A),* and *JHD2(PHDΔ)* expression was controlled by the constitutive *PYK1* promoter and delivered to the indicated yeast strains on a low copy CEN-ARS plasmid marked by *LEU2*. Each yeast strain was grown overnight in YNB-leu to maintain the plasmid. Cells were serially diluted, plated on YNB-leu plates, and grown at various temperatures. (A) *spn1-K192N* (B) *spt6-14*.

**Supp Fig. 6.**Western blot quantification.

Western blot quantification for *spt6-14* and *spt6-14 jhd2*Δ at the semi-permissive temperature (30°C) for n = 3 normalized to WT. Error bars depict 1 standard deviation. (A) H3K4me3/H3 (B) H3/Pgk1.

## References

Arigo, J. T., D. E. Eyler, K. L. Carroll and J. L. Corden, 2006 Termination of cryptic unstable transcripts is directed by yeast RNA-binding proteins Nrd1 and Nab3. Mol Cell 23: 841–851.

Belotserkovskaya, R., S. Oh, V. A. Bondarenko, G. Orphanides, V. M. Studitsky et al., 2003 FACT facilitates transcription-dependent nucleosome alteration. Science 301: 1090–1093.

Benevolenskaya, E. V., H. L. Murray, P. Branton, R. A. Young and W. G. Kaelin, Jr., 2005 Binding of pRB to the PHD protein RBP2 promotes cellular differentiation. Molecular cell 18: 623–635.

Blair, L. P., Z. Liu, R. L. Labitigan, L. Wu, D. Zheng et al., 2016 KDM5 lysine demethylases are involved in maintenance of 3’UTR length. Sci Adv 2: e1501662.

Bortvin, A., and F. Winston, 1996 Evidence that Spt6p controls chromatin structure by a direct interaction with histones. Science 272: 1473–1476.

Carroll, K. L., D. A. Pradhan, J. A. Granek, N. D. Clarke and J. L. Corden, 2004 Identification of cis elements directing termination of yeast nonpolyadenylated snoRNA transcripts. Mol Cell Biol 24: 6241–6252.

Chen, X., U. Muller, K. E. Sundling and D. A. Brow, 2014 Saccharomyces cerevisiae Sen1 as a model for the study of mutations in human Senataxin that elicit cerebellar ataxia. Genetics 198: 577–590.

Clark-Adams, C. D., D. Norris, M. A. Osley, J. S. Fassler and F. Winston, 1988 Changes in histone gene dosage alter transcription in yeast. Genes Dev 2: 150–159.

Copic, A., M. Dorrington, S. Pagant, J. Barry, M. C. Lee et al., 2009 Genomewide analysis reveals novel pathways affecting endoplasmic reticulum homeostasis, protein modification and quality control. Genetics 182: 757–769.

Costanzo, M., B. VanderSluis, E. N. Koch, A. Baryshnikova, C. Pons et al., 2016 A global genetic interaction network maps a wiring diagram of cellular function. Science 353.

Creamer, T. J., M. M. Darby, N. Jamonnak, P. Schaughency, H. Hao et al., 2011 Transcriptome-wide binding sites for components of the Saccharomyces cerevisiae non-poly(A) termination pathway: Nrd1, Nab3, and Sen1. PLoS Genet 7: e1002329.

Dai, J., E. M. Hyland, D. S. Yuan, H. Huang, J. S. Bader et al., 2008 Probing nucleosome function: a highly versatile library of synthetic histone H3 and H4 mutants. Cell 134: 1066–1078.

Darby, M. M., L. Serebreni, X. Pan, J. D. Boeke and J. L. Corden, 2012 The Saccharomyces cerevisiae Nrd1-Nab3 transcription termination pathway acts in opposition to Ras signaling and mediates response to nutrient depletion. Mol Cell Biol 32: 1762–1775.

Dehe, P. M., B. Dichtl, D. Schaft, A. Roguev, M. Pamblanco et al., 2006 Protein interactions within the Set1 complex and their roles in the regulation of histone 3 lysine 4 methylation. J Biol Chem 281: 35404–35412.

Dey, B. K., L. Stalker, A. Schnerch, M. Bhatia, J. Taylor-Papidimitriou et al., 2008 The histone demethylase KDM5b/JARID1b plays a role in cell fate decisions by blocking terminal differentiation. Molecular and cellular biology 28: 5312–5327.

Duina, A. A., 2011 Histone Chaperones Spt6 and FACT: Similarities and Differences in Modes of Action at Transcribed Genes. Genet Res Int 2011: 625210.

Formosa, T., P. Eriksson, J. Wittmeyer, J. Ginn, Y. Yu et al., 2001 Spt16-Pob3 and the HMG protein Nhp6 combine to form the nucleosome-binding factor SPN. EMBO J 20: 3506–3517.

Hainer, S. J., J. A. Pruneski, R. D. Mitchell, R. M. Monteverde and J. A. Martens, 2011a Intergenic transcription causes repression by directing nucleosome assembly. Genes & development 25: 29–40.

Hainer, S. J., J. A. Pruneski, R. D. Mitchell, R. M. Monteverde and J. A. Martens, 2011b Intergenic transcription causes repression by directing nucleosome assembly. Genes Dev 25: 29–40.

Ingvarsdottir, K., C. Edwards, M. G. Lee, J. S. Lee, D. C. Schultz et al., 2007 Histone H3 K4 demethylation during activation and attenuation of GAL1 transcription in Saccharomyces cerevisiae. Molecular and cellular biology 27: 7856–7864.

Ivanovska, I., P. E. Jacques, O. J. Rando, F. Robert and F. Winston, 2011 Control of chromatin structure by spt6: different consequences in coding and regulatory regions. Mol Cell Biol 31: 531–541.

Kaplan, C. D., L. Laprade and F. Winston, 2003 Transcription elongation factors repress transcription initiation from cryptic sites. Science 301: 1096–1099.

Kim, T., and S. Buratowski, 2009 Dimethylation of H3K4 by Set1 recruits the Set3 histone deacetylase complex to 5’ transcribed regions. Cell 137: 259–272.

Krogan, N. J., J. Dover, S. Khorrami, J. F. Greenblatt, J. Schneider et al., 2002 COMPASS, a histone H3 (Lysine 4) methyltransferase required for telomeric silencing of gene expression. J Biol Chem 277: 10753–10755.

Lenstra, T. L., J. J. Benschop, T. Kim, J. M. Schulze, N. A. Brabers et al., 2011 The specificity and topology of chromatin interaction pathways in yeast. Molecular cell 42: 536–549.

Li, Z., F. J. Vizeacoumar, S. Bahr, J. Li, J. Warringer et al., 2011 Systematic exploration of essential yeast gene function with temperature-sensitive mutants. Nat Biotechnol 29: 361–367.

Liang, G., R. J. Klose, K. E. Gardner and Y. Zhang, 2007 Yeast Jhd2p is a histone H3 Lys4 trimethyl demethylase. Nature structural & molecular biology 14: 243–245.

Lopez-Bigas, N., T. A. Kisiel, D. C. Dewaal, K. B. Holmes, T. L. Volkert et al., 2008 Genome-wide analysis of the H3K4 histone demethylase RBP2 reveals a transcriptional program controlling differentiation. Molecular cell 31: 520–530.

Lorenz, D. R., L. F. Meyer, P. J. Grady, M. M. Meyer and H. P. Cam, 2014 Heterochromatin assembly and transcriptome repression by Set1 in coordination with a class II histone deacetylase. Elife 3: e04506.

Ma, Z., D. Atencio, C. Barnes, H. DeFiglio and S. D. Hanes, 2012 Multiple roles for the Ess1 prolyl isomerase in the RNA polymerase II transcription cycle. Mol Cell Biol 32: 3594–3607.

Margaritis, T., V. Oreal, N. Brabers, L. Maestroni, A. Vitaliano-Prunier et al., 2012 Two distinct repressive mechanisms for histone 3 lysine 4 methylation through promoting 3’-end antisense transcription. PLoS Genet 8: e1002952.

Martens, J. A., L. Laprade and F. Winston, 2004 Intergenic transcription is required to repress the Saccharomyces cerevisiae SER3 gene. Nature 429: 571–574.

Martens, J. A., P. Y. Wu and F. Winston, 2005 Regulation of an intergenic transcript controls adjacent gene transcription in Saccharomyces cerevisiae. Genes & development 19: 2695–2704.

McCullough, L., Z. Connell, C. Petersen and T. Formosa, 2015 The Abundant Histone Chaperones Spt6 and FACT Collaborate to Assemble, Inspect, and Maintain Chromatin Structure in Saccharomyces cerevisiae. Genetics 201: 1031–1045.

McDonald, S. M., D. Close, H. Xin, T. Formosa and C. P. Hill, 2010 Structure and biological importance of the Spn1-Spt6 interaction, and its regulatory role in nucleosome binding. Mol Cell 40: 725–735.

Mersman, D. P., H. N. Du, I. M. Fingerman, P. F. South and S. D. Briggs, 2009 Polyubiquitination of the demethylase Jhd2 controls histone methylation and gene expression. Genes Dev 23: 951–962.

Mikheyeva, I. V., P. J. Grady, F. B. Tamburini, D. R. Lorenz and H. P. Cam, 2014 Multifaceted genome control by Set1 Dependent and Independent of H3K4 methylation and the Set1C/COMPASS complex. PLoS Genet 10: e1004740.

Nagy, P. L., J. Griesenbeck, R. D. Kornberg and M. L. Cleary, 2002 A trithorax-group complex purified from Saccharomyces cerevisiae is required for methylation of histone H3. Proc Natl Acad Sci U S A 99: 90–94.

O’Donnell, A. F., N. K. Brewster, J. Kurniawan, L. V. Minard, G. C. Johnston et al., 2004 Domain organization of the yeast histone chaperone FACT: the conserved N-terminal domain of FACT subunit Spt16 mediates recovery from replication stress. Nucleic Acids Res 32: 5894–5906.

Orphanides, G., G. LeRoy, C. H. Chang, D. S. Luse and D. Reinberg, 1998 FACT, a factor that facilitates transcript elongation through nucleosomes. Cell 92: 105–116.

Orphanides, G., W. H. Wu, W. S. Lane, M. Hampsey and D. Reinberg, 1999 The chromatin-specific transcription elongation factor FACT comprises human SPT16 and SSRP1 proteins. Nature 400: 284–288.

Porrua, O., and D. Libri, 2013 A bacterial-like mechanism for transcription termination by the Sen1p helicase in budding yeast. Nat Struct Mol Biol 20: 884–891.

Pruneski, J. A., and J. A. Martens, 2011 Transcription of intergenic DNA deposits nucleosomes on promoter to silence gene expression. Cell cycle 10: 1021–1022.

Ramakrishnan, S., S. Pokhrel, S. Palani, C. Pflueger, T. J. Parnell et al., 2016 Counteracting H3K4 methylation modulators Set1 and Jhd2 co-regulate chromatin dynamics and gene transcription. Nat Commun 7: 11949.

Santos-Rosa, H., R. Schneider, B. E. Bernstein, N. Karabetsou, A. Morillon et al., 2003 Methylation of histone H3 K4 mediates association of the Isw1p ATPase with chromatin. Molecular cell 12: 1325–1332.

Sayou, C., G. Millan-Zambrano, H. Santos-Rosa, E. Petfalski, S. Robson et al., 2017 RNA Binding by Histone Methyltransferases Set1 and Set2. Mol Cell Biol 37.

Schibler, A., E. Koutelou, J. Tomida, M. Wilson-Pham, L. Wang et al., 2016 Histone H3K4 methylation regulates deactivation of the spindle assembly checkpoint through direct binding of Mad2. Genes Dev 30: 1187–1197.

Schneider, J., A. Wood, J. S. Lee, R. Schuster, J. Dueker et al., 2005 Molecular regulation of histone H3 trimethylation by COMPASS and the regulation of gene expression. Mol Cell 19: 849–856.

Soares, L. M., M. Radman-Livaja, S. G. Lin, O. J. Rando and S. Buratowski, 2014 Feedback control of Set1 protein levels is important for proper H3K4 methylation patterns. Cell Rep 6: 961–972.

Soloveychik, M., M. Xu, O. Zaslaver, K. Lee, A. Narula et al., 2016 Mitochondrial control through nutritionally regulated global histone H3 lysine-4 demethylation. Sci Rep 6: 37942.

Steinmetz, E. J., and D. A. Brow, 1996 Repression of gene expression by an exogenous sequence element acting in concert with a heterogeneous nuclear ribonucleoprotein-like protein, Nrd1, and the putative helicase Sen1. Mol Cell Biol 16: 6993–7003.

Steinmetz, E. J., N. K. Conrad, D. A. Brow and J. L. Corden, 2001 RNA-binding protein Nrd1 directs poly(A)-independent 3’-end formation of RNA polymerase II transcripts. Nature 413: 327–331.

Takahashi, Y. H., J. S. Lee, S. K. Swanson, A. Saraf, L. Florens et al., 2009 Regulation of H3K4 trimethylation via Cps40 (Spp1) of COMPASS is monoubiquitination independent: implication for a Phe/Tyr switch by the catalytic domain of Set1. Mol Cell Biol 29: 3478–3486.

Taverna, S. D., S. Ilin, R. S. Rogers, J. C. Tanny, H. Lavender et al., 2006 Yng1 PHD finger binding to H3 trimethylated at K4 promotes NuA3 HAT activity at K14 of H3 and transcription at a subset of targeted ORFs. Molecular cell 24: 785–796.

Terzi, N., L. S. Churchman, L. Vasiljeva, J. Weissman and S. Buratowski, 2011 H3K4 trimethylation by Set1 promotes efficient termination by the Nrd1-Nab3-Sen1 pathway. Mol Cell Biol 31: 3569–3583.

Thiebaut, M., E. Kisseleva-Romanova, M. Rougemaille, J. Boulay and D. Libri, 2006 Transcription termination and nuclear degradation of cryptic unstable transcripts: a role for the nrd1-nab3 pathway in genome surveillance. Mol Cell 23: 853–864.

Tokai, M., H. Kawasaki, Y. Kikuchi and K. Ouchi, 2000 Cloning and characterization of the CSF1 gene of Saccharomyces cerevisiae, which is required for nutrient uptake at low temperature. J Bacteriol 182: 2865–2868.

Tresaugues, L., P. M. Dehe, R. Guerois, A. Rodriguez-Gil, I. Varlet et al., 2006 Structural characterization of Set1 RNA recognition motifs and their role in histone H3 lysine 4 methylation. J Mol Biol 359: 1170–1181.

Ursic, D., K. L. Himmel, K. A. Gurley, F. Webb and M. R. Culbertson, 1997 The yeast SEN1 gene is required for the processing of diverse RNA classes. Nucleic Acids Res 25: 4778–4785.

Vanacova, S., and R. Stefl, 2007 The exosome and RNA quality control in the nucleus. EMBO Rep 8: 651–657.

Vasiljeva, L., and S. Buratowski, 2006 Nrd1 interacts with the nuclear exosome for 3’ processing of RNA polymerase II transcripts. Mol Cell 21: 239–248.

Xu, M., M. Soloveychik, M. Ranger, M. Schertzberg, Z. Shah et al., 2012 Timing of transcriptional quiescence during gametogenesis is controlled by global histone H3K4 demethylation. Dev Cell 23: 1059–1071.

Zhang, K., W. Lin, J. A. Latham, G. M. Riefler, J. M. Schumacher et al., 2005 The Set1 methyltransferase opposes Ipl1 aurora kinase functions in chromosome segregation. Cell 122: 723–734.

